# Injectable Progranulin-derivative Atsttrin loaded Protein Engineered Gels for Post Traumatic Osteoarthritis

**DOI:** 10.1101/2021.05.12.443716

**Authors:** Priya Katyal, Aubryanna Hettinghouse, Michael Meleties, Changhong Chen, Min Cui, Guodong Sun, Sadaf Hasan, Rajiv Menon, Bonnie Lin, Ravinder Regatte, Chuan-ju Liu, Jin Kim Montclare

## Abstract

Protein-based biomaterials offer several advantages over synthetic materials, owing to their unique stimuli-responsive properties, biocompatibility and modular nature. We have successfully developed protein block polymers that consist of elastin like polypeptide (E) and the coiled-coil domain of cartilage oligomeric matrix protein (C). Here, we demonstrate that E_5_C, a construct consisting of five repeats of E and a single domain of C, is capable of forming a porous networked gel at physiological temperature, making it an excellent candidate for injectable biomaterials. Combination of E_5_C with Atsttrin, a chondroprotective engineered derivative of anti-inflammatory growth factor progranulin (PGRN), provides a unique biochemical and biomechanical environment to protect against post-traumatic osteoarthritis (PTOA) onset and progression. E_5_C gel was demonstrated to provide prolonged release of Atsttrin and inhibit chondrocyte catabolism while facilitating anabolic signaling *in vitro*. We also provide *in vivo* evidence that prophylactic and therapeutic application of Atsttrin-loaded E_5_C hydrogels protected against PTOA onset and progression in a rabbit anterior cruciate ligament transection model. Collectively, we have developed a unique protein-based gel capable of minimally invasive, sustained delivery of prospective therapeutics, particularly the PGRN-derivative Atsttrin, for prevention of OA onset and progression.

## Introduction

Osteoarthritis (OA) is the most prevalent chronic joint disease and is associated with substantive financial and humanistic burden [1]. Post traumatic osteoarthritis (PTOA) is a subset of OA characterized by damage to articular cartilage induced by a sudden application of mechanical force [2]. Approximately 60% of patients who suffer anterior cruciate ligament (ACL) injuries develop PTOA within 10–15 years and symptomatic OA resultant of injury is associated with $3 billion in direct U.S. health care costs [3, 4]. Neither a disease-modifying OA drug nor a non-surgical cure presently exists [5, 6]. While nonsteroidal anti-inflammatory drugs (NSAIDs) and steroids can attenuate PTOA-related pain, they have no effect on disease progression, and their use can be limited by potential severe side effects. Moreover, limited retention of soluble materials in the joint space hampers efficacy of local delivery of potential therapeutic agents. The current end-stage treatment, total joint replacement, is not desirable because of the frequent need for revision surgeries, particularly for younger individuals that develop OA prematurely as a result of a joint injury [7].

Trauma-induced cartilage injury is difficult to treat because cartilage has a limited capacity to regenerate avascular, aneural, and alymphatic tissue, and has a complex structure with unique mechanical demands [7]. Sudden impact can severely damage the structural integrity and biomechanical properties of cartilage, leading to initial responses of inflammation and apoptosis, characterized by release of cytokines, metalloproteinases, and matrix fragments [6]. This initial inflammatory phase is followed by an intermediate phase of catabolic and anabolic processes for cartilage repair [5, 6]. In the late phase, anabolic processes predominate, characterized by high levels of growth factors and matrix formation. To treat PTOA, it is important to suppress the inflammation/catabolic reaction and to induce cartilage regeneration by providing an optimal biomechanical and biochemical environment.

Progranulin (PGRN), also referred to as granulin epithelin precursor (GEP) and P-cell derived growth factor (PCDGF), is a widely expressed 593-amino acid secretory growth factor with diverse functions in numerous biological processes, including chondrocyte homeostasis [8–11]. Global screening identified PGRN as a previously unrecognized inhibitor of master inflammatory cytokine TNFα and binding partner of TNFR1/2 [12]. This PGRN/TNFR interaction and its inhibition of TNFα has been implicated in various animal models of inflammatory and degenerative disease, including arthritis [8, 9, 12–25].

Importantly, we have developed an engineered protein, Atsttrin (Antagonist of TNF/TNFR Signaling via Targeting to TNF Receptors), which comprises three TNFR-binding fragments of PGRN and is more effective than PGRN in treating inflammatory arthritis [12, 26, 27]. Similar to PGRN, Atsttrin exerts its chondroprotective effects in OA through two pathways: a) inhibition of TNFα/TNFR1 inflammatory/catabolic pathway, and b) activation of TNFR2 protective/anabolic pathway [12]. In addition, a recent study from another group demonstrated that intra-articular transplantation of Atsttrin-transduced mesenchymal stem cells ameliorated OA development [28]; however, such injections suffer from rapid diffusion with low retention of Atsttrin detected within the joint. To circumvent the issue of diffusion, Wang *et al*. utilized 3D printed Atsttrin incorporated scaffolds of alginate (Alg)/hydroxyapatite [29]. These scaffolds were capable of releasing Atsttrin over a period of seven days but did not exhibit satisfactory bone repair effects in mice. The surgical step of placing the implants is an invasive procedure, which poses some significant challenges and potential health risks. These results indicate the protective effects of Atsttrin, however, a suitable drug delivery vehicle is needed that can be easily administered, provides a sustained release of Atsstrin, and improves patient compliance and comfort.

Over the past few years, an increasing number of injectable *in situ* gel forming systems have been developed [30–34]. Such drug delivery systems exhibit phase transition from sol to gel under physiological conditions. They can be easily administered, and upon administration, they create depots under physiological temperature, providing local release of the encapsulated drug at the site of injection. Many different thermo-responsive *in situ* gels have been investigated for their use in various diseases including diabetes [30], wound healing [33], and spinal cord injuries [31]. A thermoresponsive hydrogel has been recently employed to deliver TNF inhibitor Enbrel into a PTOA mouse model through intra-articular injection [32, 34]. While there are several polymeric in situ systems reported in the literature, only a small number of protein-based systems have been proposed for joint conditions. We hypothesize that a unique combination of Atsttrin and an injectable protein engineered hydrogel would enhance the Atsttrin efficacy by increasing the retention time of Atsttrin within the arthritic joint as well as stabilizing proteins and releasing them in a sustained manner. In addition, we expect Atsttrin-loaded protein engineered hydrogel to be delivered via a single injection, avoiding the surgical procedures associated with implants.

Towards this goal, we have utilized a protein block polymer, referred to as E_5_C [35, 36], which consists of five repeats of elastin like polypeptide (ELP or E) connected to a coiled-coil domain of cartilage oligomeric matrix protein (COMPcc or C). E comprises of repeating units of pentapeptide sequence (VPGXG)_n_, where X can be any amino acid except proline and n is the number of repeating units. E can self-assemble into beta-spiral like structure and exhibits a lower critical solution temperature (LCST) behavior. C, on the other hand, has a unique characteristic to self-assemble into a homo-pentameric structure with a hydrophobic core that can bind and release small hydrophobic molecules [37]. Preliminary microrheological assessment indicated that E_5_C can self-assemble to form an elastic gel [38]. Here, we advance characterization of mechanical and rheological properties of the E_5_C gel and describe the mechanism of gelation. We demonstrate that E_5_C exhibits sol-gel behavior, existing as a solution (sol) at lower temperatures and undergoing micelle formation and crosslinking to generate a complex solid gel at physiological temperature. It is able to provide sustained release of Atsttrin over a prolonged time period. Prolonged drug release from E_5_C hydrogel, biocompatibility, and pro-anabolic and anti-catabolic effects of Atsttrin-EC hydrogel on chondrocyte metabolism are demonstrated *in vitro*.

Magnetic resonance (MR) and micro-computed tomography (µCT) imaging, cartilage macroscopic and histological scoring, and immunohistochemical staining results support *in vivo* preventive effects of Atsttrin-loaded E_5_C hydrogel on OA onset. Mild inhibition of established OA progression is indicated by macroscopic and histological scoring following therapeutic application of Atsttrin-loaded E_5_C hydrogels. Collectively, this study provides evidence supporting application of the E_5_C hydrogel, particularly Atsttrin-loaded E_5_C hydrogel as a potential therapeutic drug delivery system for OA treatment.

## Results and Discussion

### Thermoresponsiveness and self-assembling properties of E_5_C

We selected and biosynthesized E_5_C construct, which bears five repeats of E fused to C at its C-terminus (**Fig. S1**). To assess the thermoresponsiveness and assembly properties of E_5_C, we performed a series of experiments including circular dichroism (CD) spectroscopy, critical micelle temperature (CMT) and dynamic light scattering (DLS). As previously noted [36], E_5_C exhibited a random coil-like structure at lower temperatures, which transitioned to helical/beta-like conformation with increasing temperatures (**Fig. 1A**). CMT and DLS experiments indicated the ability of E_5_C to self-assemble into micellar nanoparticles (**Fig. S2, and Fig. 1B**). Nile red experiments exhibited increased fluorescence under hydrophobic conditions and was used as a probe to detect critical temperatures for micelle formation [39]. Incubation of Nile red and E_5_C was carried out at varied temperatures; fluorescence intensity gradually increased with temperature, reaching peak intensity at approximately 35 °C followed by a drop in emission intensity in the temperature range of 40 °C to 50 °C. These results indicated assembly of E_5_C protein polymer into micelles in a temperature range of 20-40 °C. At temperatures higher than 50 °C, there was a sharp rise in the fluorescence intensity indicating hydrophobic collapse as a result of aggregation. The effect of temperature on E_5_C particle size was also assessed using DLS. Z-average diameter was measured over a range of 4-65 °C (**Fig. 1B and Fig. S2B**). At 20 °C, an average size of 48.5 ± 8.0 nm was observed with particles increasing in size with an increase in temperature. Beyond physiological temperature, there was a drastic increase in the diameter with micron sized particles observed at higher temperatures.

**Figure 1.**
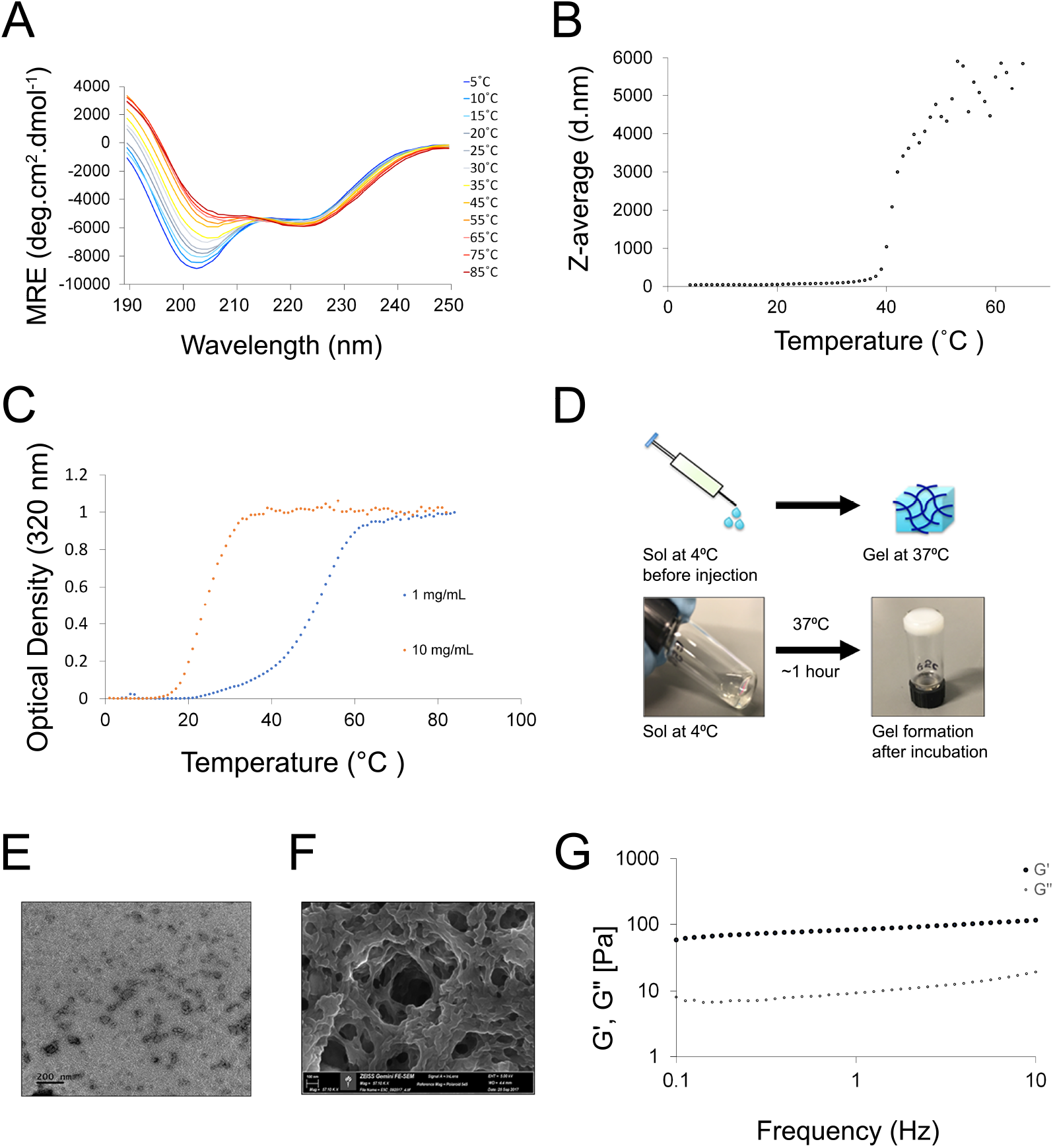
Characterization of E_5_C and gel formation. (A) Representative wavelength scans of E_5_C as a function of temperature. (B) Particle size (diameter) of E_5_C as a function of increasing temperatures as observed via DLS. (C) Optical density measurements of E_5_C under dilute (1mg/mL) and concentrated (10mg/mL) conditions. (D) Schematic of gel formation; E_5_C exists as solution (sol) at lower temperatures and solid gel at 37 °C. The approach combines syringe ability of sol and *in situ* gelling properties of a gel. Tube inversion test showing sol-gel transition using 1mM of E_5_C in 1x PBS at pH 7.4. The gel is formed within 1 hour of incubation at 37 °C. (E) Micrograph depicting micelles formation at 2 mg/mL. Scale bar represents 200 nm. (F) Scanning electron micrographs of E_5_C showing porous network formation. Scale bar represents 100 nm. (G) Rheological assessment of the storage (G′) and loss (G″) moduli of E_5_C gel at 37°C.

The transition temperature (T_t_) of E_5_C was measured via UV-Vis. The transition temperature (T_t_), defined as the temperature at which the OD value is 50% of its maximum value, varied with respect to the concentration. A higher transition temperature of approximately 50°C was observed at 1mg/mL, which was shifted to lower temperatures as the concentration was increased to 10 mg/mL (**Fig. 1C**). Turbidity profile of samples higher than 10 mg/mL did not give reliable results due to oversaturation. E_5_C exhibited an irreversible phase transition as the turbid solution did not revert back to its original state upon cooling.

### Gel formation and characterization of E_5_C network

Purified E_5_C was concentrated to approximately 1 mM (23 mg/mL) in 1x PBS, pH 7.4 and incubated in glass vials at 37°C. Gel formation was assessed using a tube-inversion test. Gelation was confirmed when a solid network was obtained that did not flow under its own weight. The E_5_C at 1 mM formed a white turbid network within an hour of incubation at 37°C (**Fig. 1D**). We hypothesized the role of micelles in forming networks that pack together to form gels. The morphologies of the micelles were further explored using transmission electron microscopy (TEM) (**Fig. 1E**). At 2 mg/mL, particles measuring 80.3 nm were observed, while at 23 mg/mL, an interconnected network of hydrogel was observed (**Fig. S3**). Scanning electron micrography (SEM) of E_5_C indicated its porous microstructure with an interconnected network (**Fig. 1F**). Oscillatory frequency sweeps were used to determine the elastic (G′) and viscous moduli (G″) as a function of frequency from 0.1-1 Hz with a strain of 1%. Measurements carried out at 37°C indicate that G′ was greater than G″ (**Fig. 1G**), confirming the elastic character of the gel.

### Extended Drug delivery via Atsttrin-loaded E_5_C Hydrogel exerts pro-anabolic and anti-catabolic effects of Atsttrin *in vitro*

An indirect ELISA using a specific anti-Atsttrin antibody was established (**Fig.2A**) in accordance with our previous publication and implemented to monitor release of Atsttrin from E_5_C hydrogel (**Fig. 2B**) [12]. Briefly, E_5_C was mixed with Atsttrin and incubated at 37 °C. The gel was washed with phosphate buffered saline to remove any free protein prior to incubating with fresh buffer. The amount of Atsttrin released into the supernatant was measured via indirect ELISA at regular intervals and the cumulative release was plotted as a function of time over a 30-day course. The E_5_C gel provided sustained release of loaded protein over prolonged time periods.

**Figure 2.**
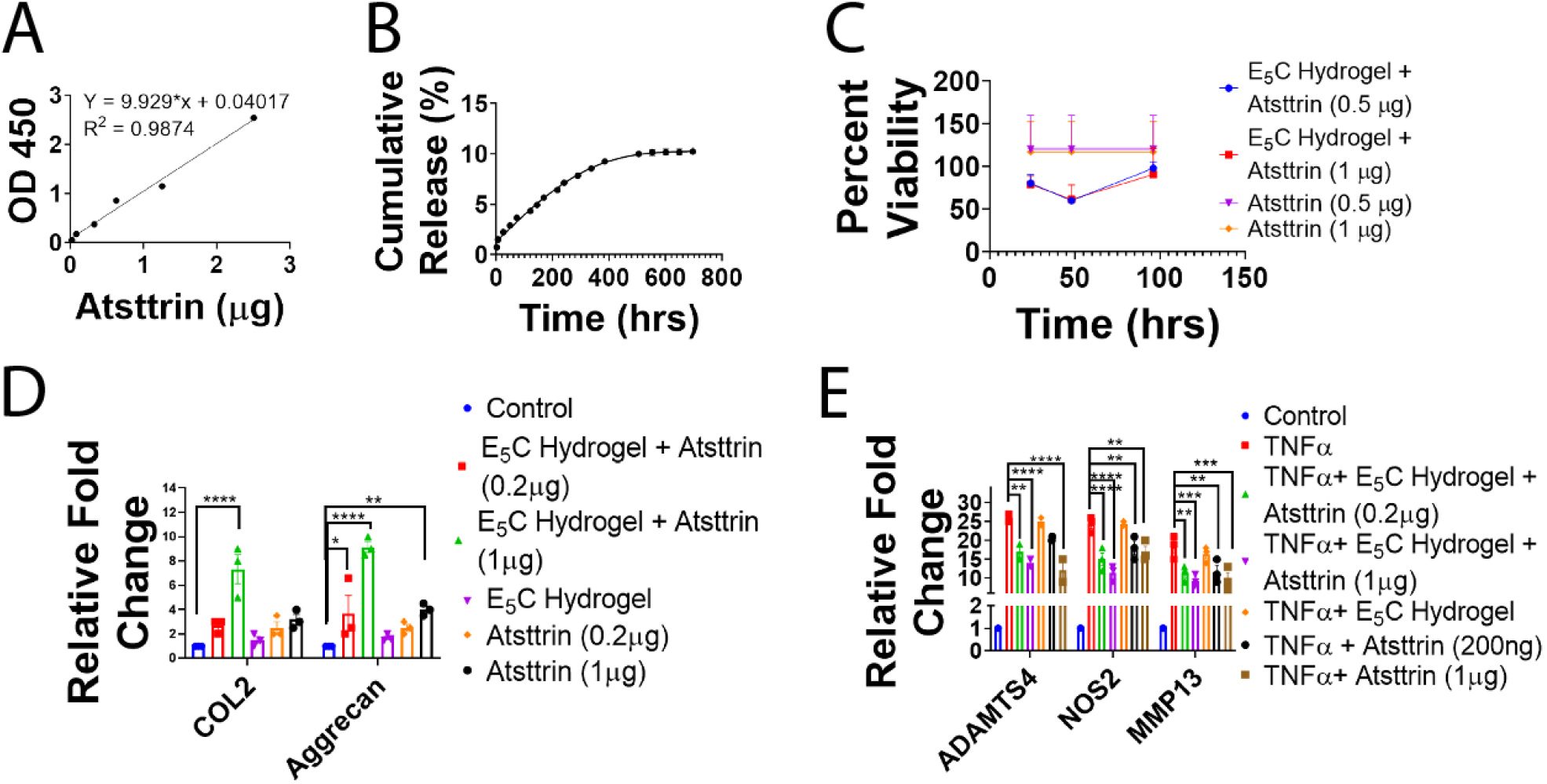
Pro-anabolic and anti-catabolic effects of Atsttrin in C28I2 chondrocytes. (A) Standard curve for indirect ELISA assay for Atsttrin. (B) Cumulative drug release curve reflecting Atsttrin release from E_5_C hydrogel over 30 days. (C) Percent viability of C28I2 cells incubated with indicated doses of Atsttrin or Atsttrin-loaded E_5_C Hydrogel. Values are normalized to appropriate control cells in which Atsttrin was omitted from culture media or hydrogel. (D) mRNA levels of anabolic markers Col2 and ACAN following treatment of C28I2 cells with Atsttrin alone or Atsttrin loaded (0.2 µg or 1 µg/mL) in hydrogels as measured by real-time PCR. (E) mRNA levels of catabolic markers ADAMTS4, NOS2 and MMP13 following stimulation of C28I2 cells with TNFα (20 ng/mL) in the presence or absence of Atsttrin alone or Atsttrin loaded (0.2 µg or 1 µg/mL) in hydrogels as measured by real-time PCR. ANOVA values presented as mean ± SEM of three independent experiments. *:p≤0.05, **:p≤0.01,***:p≤0.001, ****p<0.0001.

Absence of cytotoxic effects of Atsttrin and E_5_C hydrogel loaded with Atsttrin (0.5 or 1 µg/mL) was also confirmed using the tetrazolium (MTT)-based colorimetric assay (**Fig. 2C**). To study the effect of Atsttrin-loaded E_5_C hydrogel on chondrocyte metabolism, endotoxin-free E_5_C and Attstrin or sterile phosphate buffered saline (PBS) were mixed at a ratio of 10:1 v/v and 200 µL of the resulting mixture was coated onto the surface of an 8.0 µm pore size polycarbonate cell culture insert and incubated at 37 °C. Following gel formation, inserts were introduced to 24-well plates containing C28I2 chondrocyte cells and cells were assayed for anabolic and TNF-α induced catabolic response genes by real time polymerase chain reaction (PCR) following 48 hour culture. Atsttrin alone and E_5_C hydrogel loaded with Attstrin stimulated dose dependent inhibition and stimulation of catabolic (matrix metalloproteinases (MMP13), NOS2, and ADAMTS4) and anabolic markers (Collagen II (Col2), Aggrecan), respectively, at the mRNA level *in vitro* (**Fig. 2D,E**). Moreover, data reflected a dose dependent effect of Atsttrin and E_5_C hydrogel loaded with Atsttrin on inhibition of catabolic and stimulation of anabolic markers at the mRNA level (**Fig. 2D,E**).

### *In vivo* gelation and intra-articular retention of the hydrogel

Adult DBA1/J mice were randomly divided into groups receiving either Vivo-Tag 680-S (NEV10121, Perkin Elmer) labeled E_5_C, Vivo-Tag-S 680 labeled Atsttrin (5µg) diluted in E_5_C, or Vivo-tag-S 680 labeled Atsttrin (5µg) diluted in PBS via a 5 µL bilateral intra-articular injection. The intra-articular retention of either hydrogel or Atsttrin was monitored via total fluorescence intensity using an IVIS Spectrum imaging system (PerkinElmer, Fremont, USA) at days 0, 2, 4, 14, 21, 28 and 32 after injection. The total fluorescence intensity was normalized against background signal and relative fluorescence intensity (RFI) was plotted over time (**Fig. 3**). The signal had reached zero in all groups by 32 days post-injection. Importantly, fluorescence signal was more robustly retained in groups that received E_5_C hydrogel or Atsttrin-loaded E_5_C, indicating that E_5_C was capable of gelation and prolonging retention of loaded biologics *in vivo* (**Fig. 3**).

**Figure 3.**
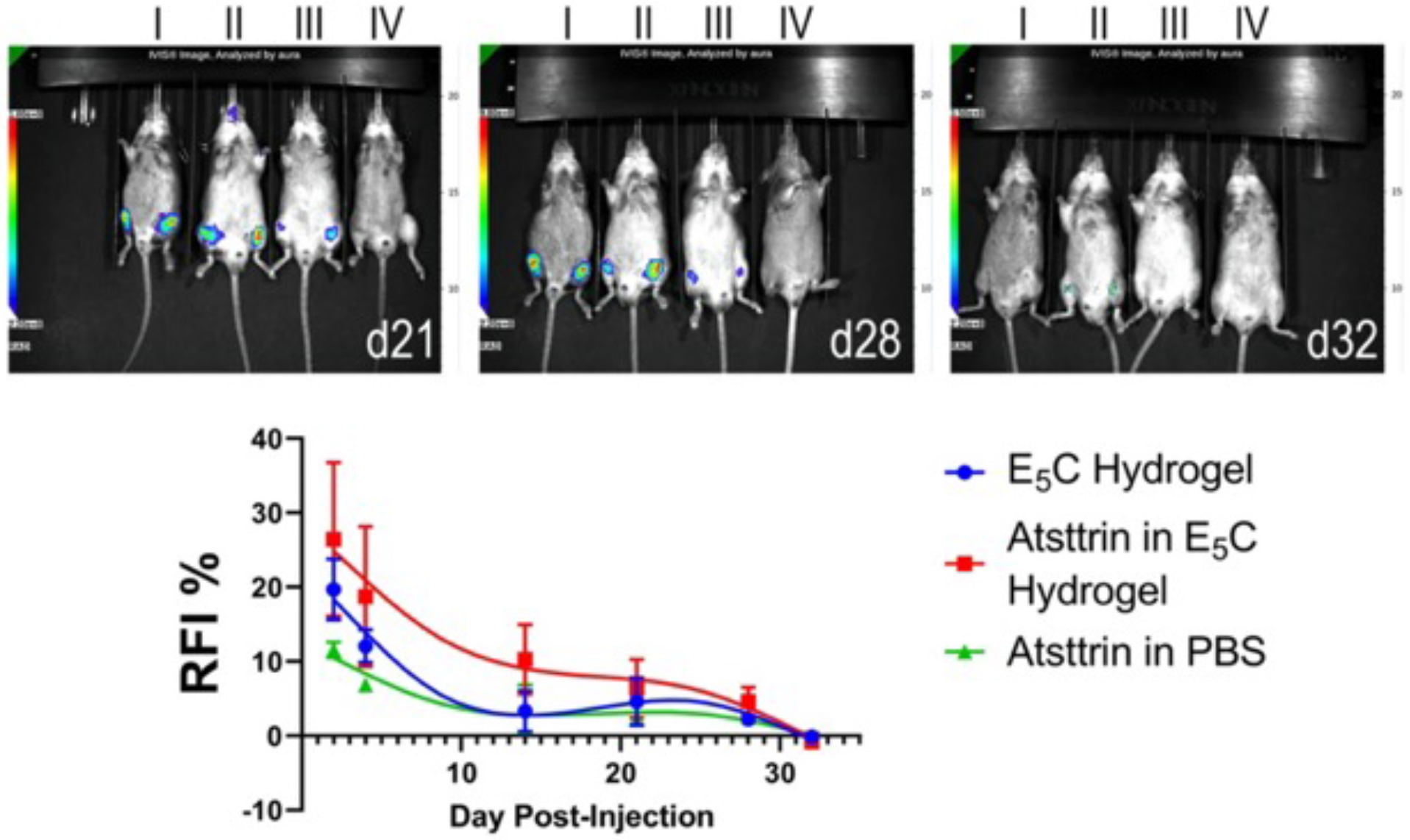
IVIS imaging of localized delivery of the E_5_ C hydrogel. Mice (n=2/group) received bilateral intra-articular injections of phosphate buffered saline or fluorescently labeled E_5_C or Atsttrin; I: Vivo-Tag 680 labeled E_5_C, II: Vivo-Tag 680 labeled Atsttrin in E_5_C, III: Vivo-Tag 680 labeled Atsttrin in phosphate buffered saline, IV: phosphate buffered saline. As shown, signal remains localized to the knee joint at each timepoint; the control mouse shows no positive signal in the joint. In groups receiving E_5_C, lack of diffusion beyond the joint area indicates successful gelation and localized retention of loaded Atsttrin.

### Atsttrin-loaded E_5_C Hydrogel protects cartilage quality during OA onset and progression

To assess the preventative potential of our drug and drug loaded E_5_C hydrogels *in vivo*, we established a PTOA model in mature male New Zealand rabbits (Charles River Laboratories) via bilateral anterior cruciate ligament transection (ACLT, n=36) or a sham operation (n=6). Prospective biologics were delivered with a single intra-operative intra-articular injection of 0.5 mL E_5_C alone (n=6), Atsttrin alone (0.5 mg; n=6), or E_5_C loaded with Atsttrin (0.5mg; n=6), or vehicle (sterile phosphate buffered saline; n=12) (**Fig.4A**). Animals were permitted to develop a PTOA phenotype over 8 weeks prior to sacrifice [40].

**Figure 4.**
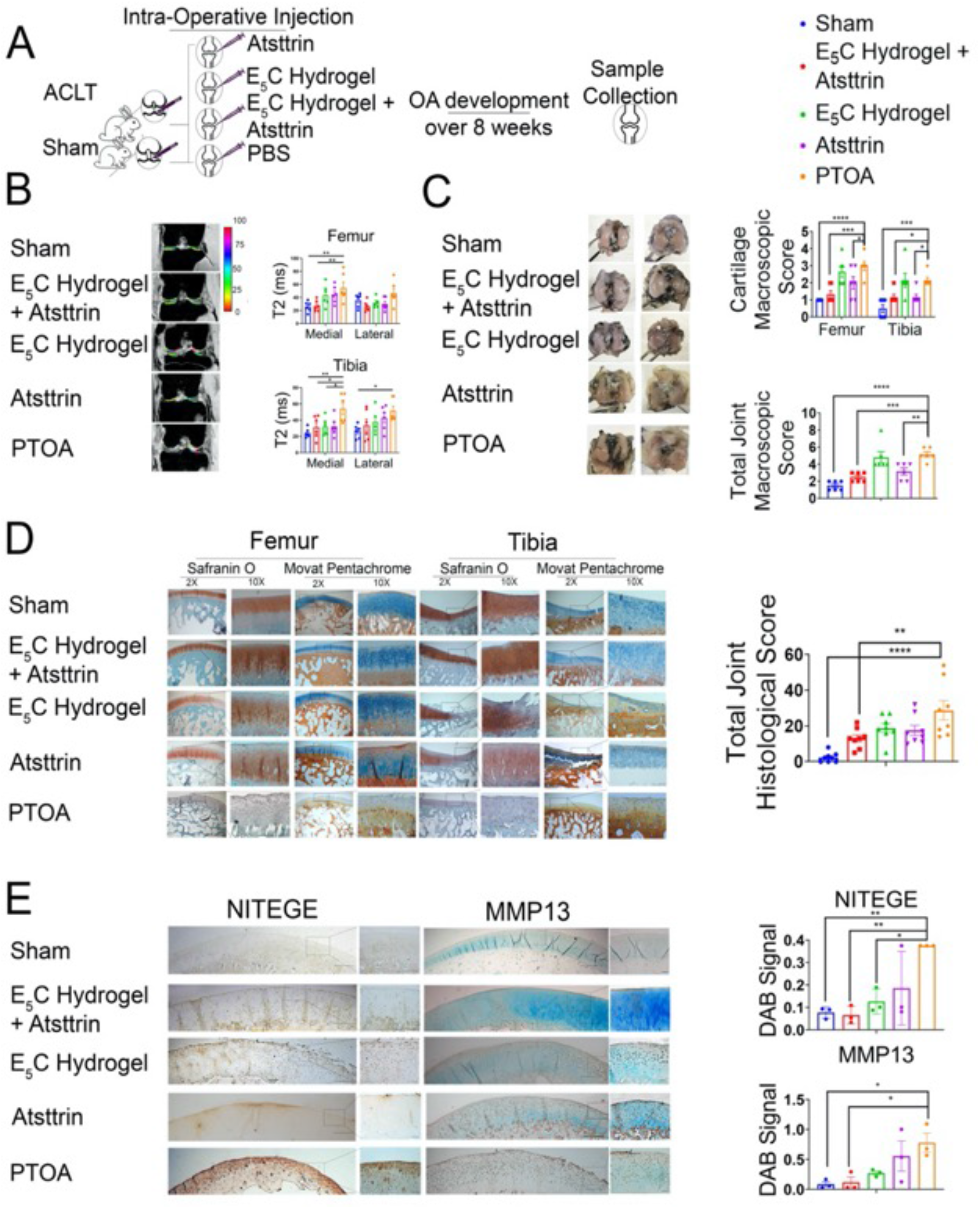
Atsttrin-loaded E_5_C hydrogel protects cartilage quality during OA onset. (A) Schematic illustrating PTOA induction for evaluation of preventative potential of indicated treatments. (B) Representative T2 map overlay images and quantification of T2 extinction times (ms) for cartilage of the lateral and medial compartments of the femur and tibia. n=6/group. (C). Representative images of India-Ink stained femoral heads and tibial plateaus. Quantitation of macroscopic score, n=6/group. (D) Representative images from Safranin O / Fast Green staining of femoral condyles and tibial plateaus, quantification of cumulative histological score, n=8/group. (E). Immunohistochemical staining for NITEGE and MMP13. Antibody staining shown by development of 3,3′-diaminobenzidine (DAB) signal (brown) followed by counterstaining with methyl green. Signal quantified relative to nuclei in ImageJ. n=3/group. ANOVA values presented as mean ± SEM. *:p≤0.05, **:p≤0.01, ***: p≤0.001. Scale bars are 100 μm.

Following sacrifice, left hind limbs were collected for magnetic resonance (MR) imaging. Quantification of cartilage spin-spin relaxation (T2) values demonstrated significantly lower T2 values in regions of interest highlighting weight bearing cartilage of the medial femoral condyle and medial tibial plateau of sham animals and those treated intraoperatively with Atsttrin-loaded E_5_C hydrogel relative to those from animals receiving no preventative treatment correlating to maintenance of cartilage integrity[41] (**Fig. 4B**). In femurs and tibias, T2 values were not significantly different between PTOA and Atsttrin treatment groups, while rabbits that had received E_5_C alone did show some benefit in the medial tibia relative to PTOA animals (**Fig. 4B**). Following MR scanning, limbs were disjointed and stained with India Ink for macroscopic assessment of cartilage [42]. Deposition of India ink within areas of cartilage damage at femoral heads and tibial plateaus (**Fig. 4C)**, revealed noticeably greater fibrillation in PTOA rabbits as well as those treated with E_5_C hydrogel or Atsttrin alone relative to sham operated animals and Atsttrin-loaded E_5_C hydrogel treated animals. Surprisingly, animals receiving Atsttrin alone also exhibited statistically significant reduction in macroscopic scores relative to PTOA animals (**Fig. 4C)**.

After routine tissue processing, histological stains were applied to further evaluate cartilage quality at femoral heads and tibial plateaus. Representative images of safranin O and movat pentachrome-stained medial femoral heads and tibial plateaus are shown in **Fig. 4D**. Medial and lateral aspects of the joint were subjected to semi-quantitative grading of articular cartilage and summed to generate whole limb total scores [42]. Total scores for each limb indicated a significant chondroprotective effect of the Atsttrin-loaded E_5_C hydrogel (**Fig. 4D**). Though histological scores were reduced for animals that had received E_5_C or Atsttrin relative to those associated with untreated PTOA, these results did not reach statistical significance (**Fig. 4D**).

To explore the anti-catabolic activities of E_5_C hydrogel delivered Atsttrin, immunohistochemical staining for markers of chondrocyte catabolism including aggrecan neoepitope (NITEGE) and MMP13 was performed (**Fig. 4E**). Over the course of osteoarthritis, aggrecan is cleaved by matrix metalloproteinases (MMPs) to generate aggrecan neoepitope and elevated levels of MMP-13 are associated with OA cartilage [43, 44]. Significantly reduced positive signal for aggrecan neoepitope and reduced staining for MMP13 was observed in cartilage tissues from animals that received Atsttrin-loaded E_5_C hydrogels relative to PTOA controls (**Fig. 4 E**).

Observation of the chondroprotective effects of preventative application of our Atsttrin-loaded E_5_C hydrogels led us to examine their therapeutic efficacy in rabbit PTOA model. For this purpose, rabbits were subject to bilateral ACLT and subsequently received bilateral intra-articular injection of 0.5 mL E_5_C alone (n=3), Atsttrin alone (0.5 mg; n=3), or E_5_C loaded with Atsttrin (0.5 mg; n=3), or vehicle (sterile phosphate buffered saline; n=3) at 8 weeks post-op (**Fig. 5A**). Animals were sacrificed for analysis at 16 weeks post-op and joint tissues were collected for processing. As shown in **Fig. 5B**, macroscopic scores for femoral cartilage were significantly lower for E_5_C loaded with Atsttrin and Atsttrin treated animals than for vehicle treated PTOA animals. Similarly, histological scoring of joint tissues uncovered a localized inhibition of cartilage damage with receipt of Atsttrin-loaded E_5_C within the medial compartment of the femoral head (**Fig. 5C**)

**Figure 5.**
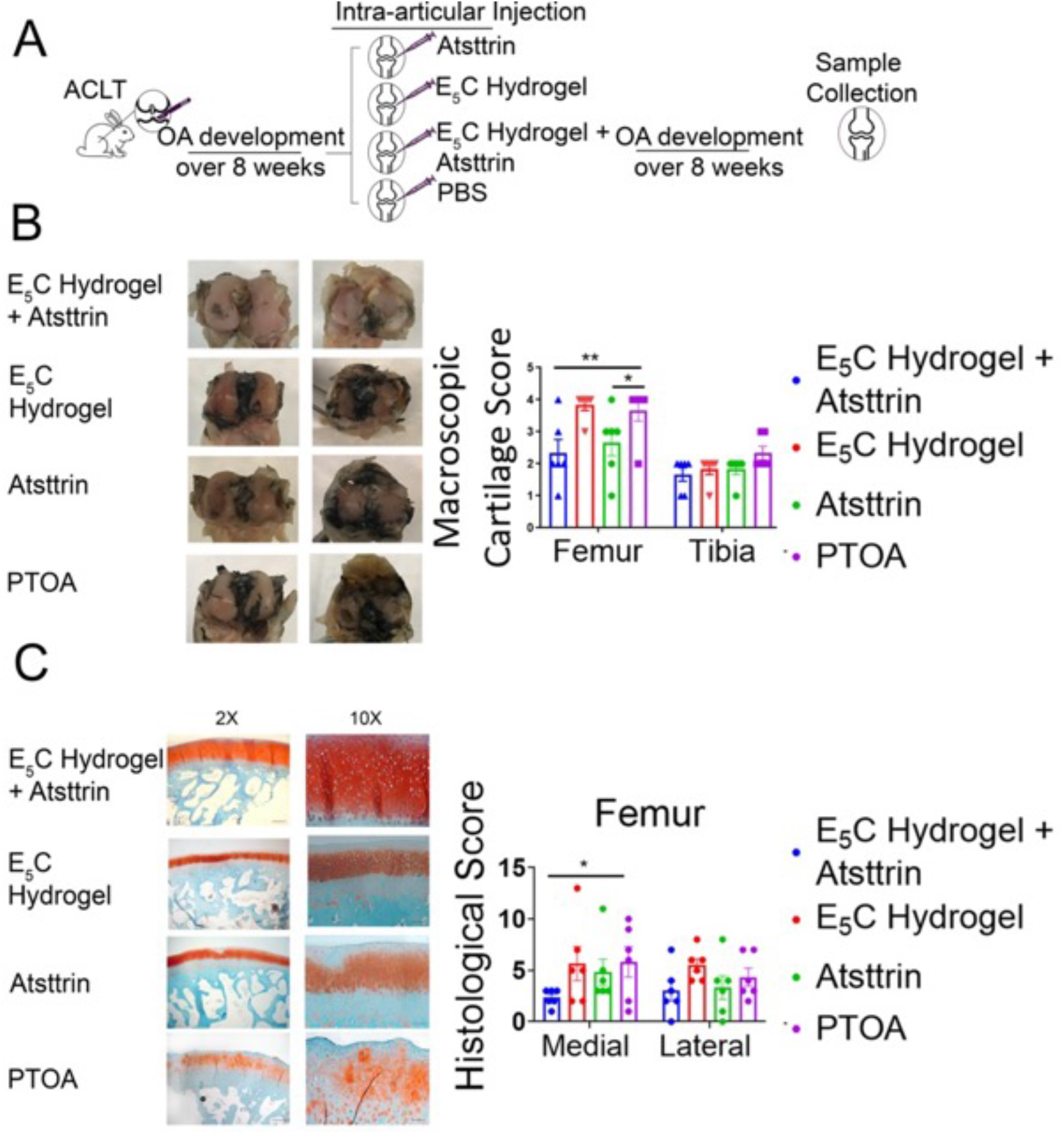
Atsttrin-loaded E_5_C hydrogel protects cartilage quality during OA progression. (A) Schematic illustrating PTOA induction for evaluation of therapeutic potential of indicated treatments. (B) Representative images of India-ink stained femoral heads and tibial plateaus. Quantitation of macroscopic score. n=6/group. (C) Representative images from Safranin O / Fast Green staining of femoral condyles, quantification of histological score. n=6/group. ANOVA values presented as mean ± SEM. *:p≤0.05, **:p≤0.01, ***: p≤0.001. Scale bars are 100 μm.

Collectively, these results indicate that E_5_C and Atsttrin exert protective effects against cartilage damage. Specifically, Atsttrin delivered via E_5_C gels has a significant preventative effect on cartilage degeneration associated with PTOA *in vivo*. Moreover, immunostaining indicates that this protective effect is due, at least in part, to inhibition of cartilage catabolism. In therapeutic application, Atsttrin delivered via E_5_C gels also exerts a localized protective effect within the medial femur.

### Atsttrin-loaded E_5_C Hydrogel protects bone quality during OA progression

microCT scanning was performed on right fixed femurs from animals in our preventative study to assess changes in bone quality. Measures of bone morphometry were analyzed in anterior, central, and posterior aspects of the weight-bearing region of femoral condyles (**Fig. 6A-H**) [45]. Bone volume fraction (BV/TV), trabecular thickness (Tb.th), trabecular number (Tb.N), and trabecular spacing (Tb.S) were analyzed in each region. Notably, animals that received intra-operative injection of E_5_C loaded with Atsttrin exhibited significantly greater BV/TV in the medial posterior and lateral central regions of interest relative to PTOA-control animals (**Fig. 6 A,E**). Tb.N and Tb.S were also significantly elevated relative to PTOA-controls in the central region of the lateral femur (**Fig. 6 G, H**) though additional measures of bone quality did not exhibit significant differences between treated and non-treated ACLT groups in the medial condyle. Averaging over treatment group without regard to laterality or subregion reveals statistically significant differences in BV/TV, Tb.N, and Tb.S between Atsttrin-loaded E_5_C hydrogel treated animals and non-treated counterparts (**Fig. 6I-L**). Tb.S was also significantly lower in animals that were treated intraoperatively with hydrogel alone, suggesting that our E_5_C construct may offer some protection against PTOA-associated changes in microstructural indices of bone **(Fig. 6L)**. Reduction of BV/TV in ACLT animals, relative to sham controls, suggested cancellous bone loss during OA development. Accordingly, BMD was assessed; averaging over treatment group revealed statistically significant differences in BMD between Atsttrin-loaded E_5_C hydrogel treated animals and non-treated counterparts (**Fig. 6M**).

**Figure 6.**
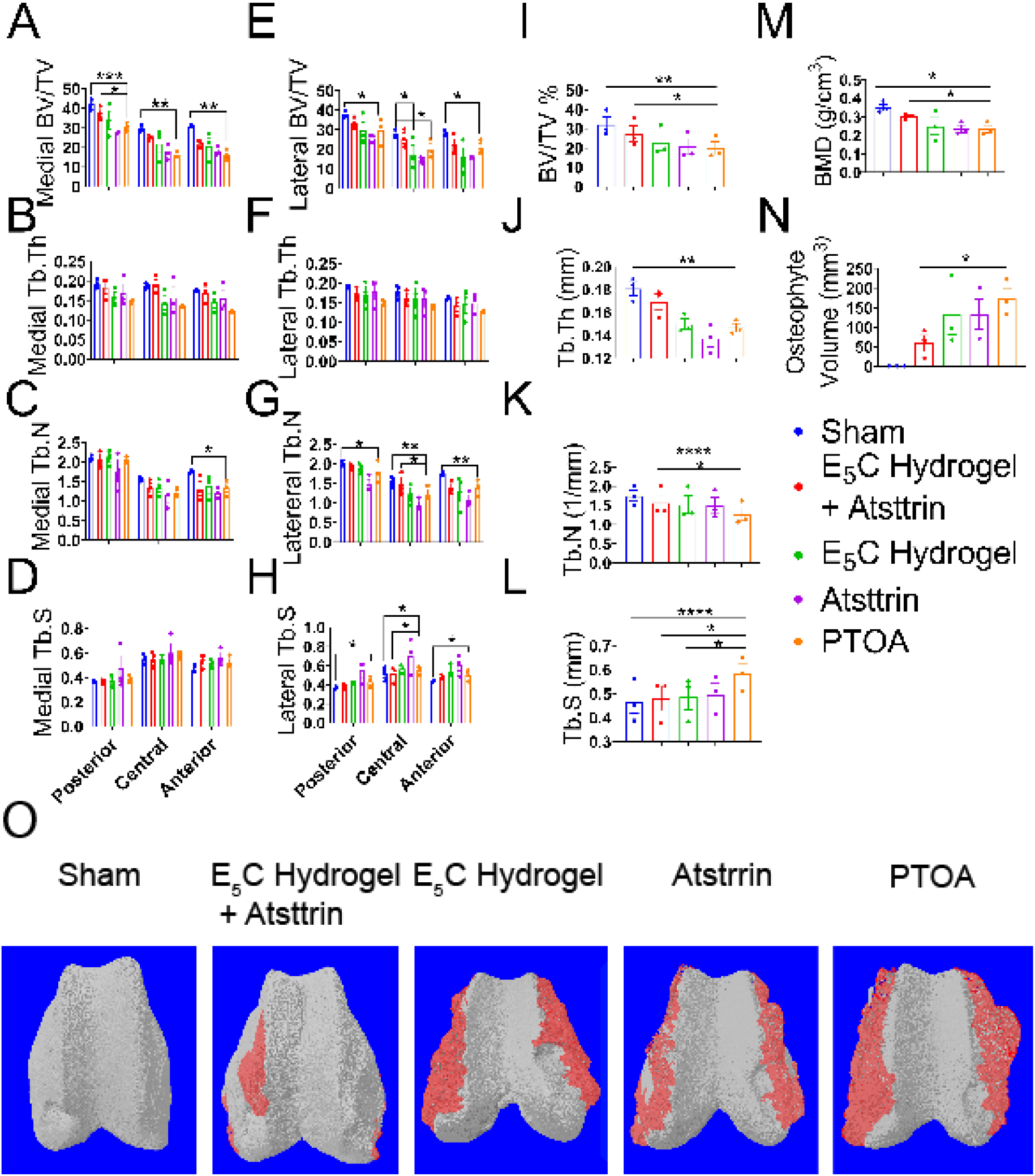
Atsttrin-loaded E_5_C hydrogel protects bone quality during OA onset. Quantification of bone morphometry within anterior, central and posterior weight bearing regions of medial (A-D) and lateral (E-H) femoral condyles. (I-L) Bone morphometric parameters regardless of laterality or subregion. (M) Quantification of bone mineral density. (N) Quantification of osteophyte volume. (O) Representative volume of interest model overlays displaying osteophytic tissue (red) and underlying bone (grey). ANOVA values presented as mean ± SEM. n=3/group, *:p≤0.05, **:p≤0.01, ***: p≤0.001

The occurrence of osteophytes is common in OA and may contribute to discomfort and loss of joint function [46]. Sham operated animals exhibited no osteophyte development while osteophyte formation occurred to varying degrees in PTOA animals. Osteophyte volume was quantified by manual segmentation of osteophytic tissue based on deviation from the appearance of the underlying normal bone contour; manual segmentation was performed every 5 slices with adaptive interpolation over the entirety of the image sequence (approximately 10 mm). The total 3D tissue volume of this region of interest was used to quantify osteophyte volume, revealing significantly less osteophytic tissue in animals treated with Atsttrin-loaded E_5_C hydrogel relative to OA-controls (**Fig. 6N**). Illustrative 3D volume models were generated in CtVox (Bruker Corporation) (**Fig. 6O**). Cumulatively, these results indicated that Atsttrin delivered via E_5_C exerts a protective effect against PTOA associated changes in bone morphometry and osteophyte development.

### Conclusions

In light of the limitations and pitfalls of available PTOA interventions, there has been growing interest in development of new drug delivery systems that are capable of providing minimally invasive, extended drug delivery [47, 48]. To this end, we investigate the ability of E_5_C to undergo smart solution-to-gel transition at body temperature. E_5_C is biocompatible, with no cytotoxic effects on human chondrocyte cells, and is biodegraded in the joint within 8-weeks post-injection. These hydrogels effectively provide a sustained and controlled release of PGRN derivative Atsttrin, a therapeutic protein known to have anti-inflammatory and chondroprotective effects. Application of Atsttrin-loaded E_5_C hydrogels exerts protective effects on cartilage and bone quality as well as therapeutic effects reflected by improved maintenance of cartilage integrity in a rabbit PTOA model. Since the therapeutic efficacy of Atsttrin-loaded E_5_C hydrogels is mild, a follow-up study is underway to optimize appropriate effective dosing. In this study, we have employed the same single drug concentration and single injection treatment strategy for both preventative and therapeutic study designs. Future exploration of higher drug loads and/or repeated drug administration in larger cohorts will allow us to optimize the utility of our Atsttrin-loaded construct in PTOA progression. Importantly, development and implementation of drug delivery systems capable of providing sustained release is desirable to provide enhanced therapeutic efficacy by maximizing local exposure to therapeutic agents and minimizing the need for repeated intra-articular injections. The current study not only supplies additional evidence supporting the protective application of Atsttrin in the pathogenesis of PTOA but also describes development of a new minimally invasive drug delivery system that may be implemented to prevent and treat PTOA and other degenerative joint diseases.

## Materials & Methods

### Materials

Pierce bicinchoninic acid (BCA) assay kit, Pierce snakeskin dialysis tubing 3.5 kDa molecular weight cut off (MWCO), Pierce high-capacity endotoxin removal spin columns, HisPur Ni-NTA agarose, Trizol®, SYBR Green Master Mix, and sodium dodecyl sulfate (SDS) were acquired from Thermo Fisher Scientific. Imidazole was purchased from Acros Organics. Acrylamide/bis solution (30%) 29:1 and natural polypeptide sodium dodecyl sulfate-polyacrylamide gel electrophoresis (SDS-PAGE) standard were purchased from Bio-Rad. Macrosep and Microsep Advance centrifugal devices 3 kDa MWCO and 0.2 μm syringe filters were purchased from Pall Corporation. Formvar/carbon-coated copper grids and uranyl acetate (1% solution) for TEM were purchased from Electron Microscopy Sciences. Silicon wafers for scanning electron microscopy (SEM) were purchased from Ted Pella, Inc. VivoTag-S 680 was purchased from PerkinElmer. Amicon® Ultra-15 Centrifugal Filter Units (3 kDa MWCO) were purchased from Millipore Sigma. Atsttrin was supplied by Synermore Biologics Co., LTD. Anti-Atsttrin antibody was supplied by Lampire Biological Laboratories (Pipersville, PA, USA). Biotinylated goat Anti-Rabbit IgG Antibody (H+L), biotinylated goat Anti-Mouse IgG (H+L) and VECTASTAIN® ABC-HRP Kit were purchased from Vector Laboratories. Anti-Aggrecan Neoeptiope (NITGE) antibody was purchased from Mybiosource. Anti-MMP13 antibody was purchased from ThermoFisher. Penicillin-Streptomycin solution, Safranin O stain, Fast Green stain, and Thiazolyl Blue Tetrazolium Bromide (MTT) and were purchased from Sigma-Aldrich. High-Capacity cDNA Reverse Transcription Kit was purchased from Applied Biosystems. Movat Pentachrome staining kit was purchased from Abcam.

### E_5_C construct Expression and Purification

E_5_C (cloned in pQE30 vector) was transformed in phenylalanine auxotrophic *E. coli* strain (AFIQ) cells and plated on tryptic soy agar plate with 0.2 mg/mL ampicillin and 0.35 mg/mL chloramphenicol. Colonies were allowed to grow at 37 °C for 12-16 hours. A single colony was inoculated in 5 mL minimal media (0.5 M Na_2_HPO_4_, 0.22 M KH_2_PO_4_, 0.08 M NaCl, 0.18 M NH_4_Cl, 1g/L of 20 amino acids, 20% w/vol. glucose, 0.1 mM CaCl_2_, 1 mM MgSO_4_, and 0.35 mg/mL vitamin B1) supplemented with 0.2 mg/mL ampicillin and 0.35 mg/mL chloramphenicol, and grown overnight. A 50 mL secondary culture was seeded with the 5 mL starter culture and allowed to grow for 2 hours at 37 °C. Approximately, 25 mL of secondary culture was transferred to 200 mL fresh minimal media, and induced with autoinduction media (0.5% glycerol, 0.05% glucose and 0.2 % lactose). The culture was grown at 37 °C for 7.5-8 hours. The cells were harvested at 4 °C and 4000 rpm for 25 minutes. The cells were re-suspended in lysis buffer (50 mM sodium phosphate dibasic pH 8, 6 M urea, and 20 mM imidazole). Purification was performed under denaturing conditions using a probe sonicator (Qsonica Q500) at 60% amplitude for 2 minutes (5 seconds on, 30 seconds off). The lysate was centrifuged at 14000 rpm at 4 °C for 50 mins to remove insoluble cell debris. The supernatant was incubated with 2.5 mL of prepared nickel beads at 4 °C for overnight. The mixture was transferred to a gravity-column and the flow-through was collected. The beads were washed with 35 mL lysis buffer, and the protein was eluted using a concentration gradient of imidazole ranging from 25 mM to 500 mM. The eluates were analyzed using (SDS-PAGE (**Fig. S1**) and the fractions containing E_5_C were pooled and dialyzed. Imidazole and urea were removed via bucket dialysis over 6x5L of 1x PBS, pH 7.4. The dialyzed protein was concentrated using 3 kD MWCO Macrosep and Microsep Advance centrifugal filters at 4 °C. Protein concentration was measured using Pierce BCA assay kit.

### Circular Dichroism

Circular dichroism (CD) was performed using Jasco J-815 CD spectrometer (*Jasco, MD, USA*) equipped with a PTC-423S single position Peltier temperature control system. Wavelength scans were performed on 4 μM protein samples in 1x PBS, pH 7.4 over the range of 200−250 nm with a bandwidth of 1 nm and path length of 1 cm. Temperature scans were performed at 204 nm from 4 to 85 °C, at a heating rate of 1°C/min. Data analysis was performed as reported by Hill *et. al*[49].

### Critical Micelle Temperature (CMT)

15 µL of 1 μM Nile red dissolved in methanol was added to PCR tubes and dried in a vacuum chamber for at least 2 hours. 150 µL of protein solution at 0.5 mg/mL were added to each tube at 4 °C with gentle mixing. The tubes were transferred to a thermocycler and slowly heated from 4-65°C with 10 min incubation periods at each measured temperature. At a given temperature, 100 µL of solution was pipetted into a 96-well solid black, round bottom polystyrene microplate and subjected to fluorescence measurements. The samples were excited at 550 nm with an emission range of 575-700 nm. The peak emission occurred at 625 nm and the average fluorescence intensities from each temperature were plotted as a function of temperature.

### Dynamic Light Scattering

Particle size as a function of temperature was measured using Zetasizer Nano Series ZS90 (Malvern Instruments, Worcestershire, UK) with the following settings: material protein with a refractive index 1.450 and absorption 0.001, dispersant PBS with a viscosity of 0.8882 cP, and refractive index of 1.330. 1mg/mL E_5_C in PBS, pH 7.4 was filtered directly into the low volume cuvette using 0.2 μm syringe filter and covered with 30 μL mineral oil. The cuvette was sealed using a parafilm and the Z-average diameter was measured over 4-65°C.

### UV-Vis Spectroscopy

The transition temperature (T_t_) of E_5_C at varying concentrations of were measured using a UV-Vis Cary-50 (Varian Inc.) equipped with a TC125 temperature regulator (Quantum Northwest). The change in optical density (OD) was recorded at 320 nm as the temperature was increased from 4 to 55°C at a heating rate of 1°C/min. The values at OD_320_ were plotted as a function of temperature. The transition temperature was calculated as described previously.[49, 50]

### Gelation

A 150 µL solution of E_5_C in 1xPBS, pH 7.4 placed in a 2 mL glass vial was incubated at 37 °C at a concentration of 1 mM. Gel formation was assessed using a tube-inversion test and visually monitored over a period of 24 hours. Gelation was confirmed when solid network was observed that did not flow upon inverting the tube. Phase separation was considered when two distinct layers (clear supernatant and turbid dense phase) were observed.

### Transmission Electron Microscopy (TEM)

Approximately 2 µL of 0.1 mM and 1 mM of E_5_C sample prepared in 1xPBS, pH 7.4 were spotted on copper grids. After 1 minute, the grids were blotted using filter water and washed twice using 5 µL water (dH_2_O). The samples were negatively stained with 3 µL of 1% uranyl acetate and dried at room temperature for 10 minutes. TEM was performed using Talos L120C TEM equipped (ThermoFisher Scientific, USA) with Gatan 4k x 4k OneView camera. The particle diameters were measured using ImageJ software.[51]

### Scanning Electron Microscopy

MERLIN field emission scanning electron microscope (Carl Zeiss AG, USA) was used to study the topography of E_5_C gels. 2 µL of 1 mM E_5_C samples in 1x PBS, pH 7.4 were spotted on silicon wafers and dried at 37°C. Dried samples were iridium-coated to 3 nm using a high-resolution sputter coater 208HR (Cressington Scientific Instruments) prior to image acquisition.

### Rheology

Oscillatory shear rheometry was used to assess the mechanical properties and damping characteristics of E_5_C. The mechanical properties of the protein hydrogel were determined using a stress-controlled ARES-G2 rheometer (TA Instruments, New Castle, DE). Protein was prepared at a concentration of 1 mM in 1x PBS, pH 7.4 at 4°C. The rheometer was equipped with 20 mm diameter parallel plates and a geometry gap of 0.5 mm was used. Protein solution was applied to the instrument at 37°C, where it was allowed to gel for an hour using a temperature controller. An oscillatory strain sweep from 0.1 to 1000% strain at a frequency of 1 Hz was used to determine the linear viscoelastic region (LVE) of the gel and to characterize the gel’s fracture. Oscillatory frequency sweeps were performed in triplicate to determine the elastic modulus (G’) and viscous modulus (G’’) as a function of frequency from 0.1 to 10 Hz with a strain of 1%.

### Endotoxin removal

Pierce High-Capacity Endotoxin Removal spin columns were prepared and washed as per manufacturer’s protocol. The columns were then equilibrated with endotoxin-free 1× PBS, pH 7.4. The protein sample at a concentration of 5 mg/mL was added to the resin and incubated for 1 hour at 4 °C with end-over-end mixing (Labquake Rotisserie, Barnstead Thermolyne). The sample was eluted in a sterile tube via centrifugation at 500 g for 1 min. Protein was further concentrated to a desired concentration using Macrosep centrifugal devices.

### Atsttrin Purification

Atsttrin was supplied by Synermore Biologics Co., LTD. *E. coli* DE3 expressing pGEX-Atsttrin was employed for affinity purification using glutathione-agarose beads as previously reported [52]. Atsttrin was freed from the GST-tag via a 6 hour Factor Xa digestion. Following digestion, supernatant was collected and Factor Xa was removed with Factor Xa Removal Resin (QIAGEN). Endotoxin was removed using endotoxin removal resin and Atsttrin purity was determined by SDS-PAGE and reverse phase high performance liquid chromatography (HPLC).

### Indirect ELISA for release studies

E_5_C was mixed with Atsttrin (6 µg) and incubated at 37 °C. The gel was washed with phosphate buffered saline to remove any free protein prior to incubating with fresh buffer. At different timepoints, the entire supernatant was removed for measuring released Atsttrin. Fresh buffer was supplemented following each removal. Equal volumes of collected supernatants were coated onto the surface of a high-binding plate overnight at 4°C after which the plate was washed three times with phosphate buffered saline (PBS). The plate was blocked in 5% non-fat milk for 2 hrs at room temperature before washing twice with PBS, 100 µL of anti-Atsttrin antibody (1µg/mL) was added to each well and incubated overnight at 4°C. The plate was washed 4 times with PBS and 100 µL secondary biotinylated goat Anti-Rabbit IgG Antibody (H+L) (1:50000) was added to each well for 2 hrs at room temperature. The plate was washed 4 times with PBS prior to addition of 100 µL TMB (3,3’,5,5’-tetramethylbenzidine) solution. Plates were incubated for 15 minutes at room temperature; the reaction was stopped by addition of 100 µL H_2_SO_4_ and read at 450 nm using a spectrophotometer.

### Cell Culture

C28I2 cells were maintained as a monolayer in Dulbecco’s modified Eagle’s medium (DMEM) supplemented with 10% fetal bovine serum, 50 U/mL penicillin and 0.05 mg/mL streptomycin. To study the effect of Atsttrin-loaded E_5_C hydrogels on chondrocyte metabolism, endotoxin-free E_5_C and Attstrin were mixed at a ratio of 10:1 v/v. 200 µL of the resulting mixture was transferred to transwell inserts with an 8 µm pore size. After gel formation over a 1 hour incubation at 37 °C, transwells were inserted into 24-well plates containing C28I2 cells. Cells were assayed for anabolic and TNF-α (20ng/mL)-induced catabolic response genes by real time PCR following a 48 hour culture.

### MTT Assay

Approximately 50 µL of E_5_C, with or without Atsttrin at indicated doses, was deposited in a sterile cell culture plate and allowed to solidify over 24-48hrs. C28I2 chondrocyte cells were seeded onto the surface of hydrogels. Alternatively, Atsttrin was added to C28I2 culture media at indicated doses. At each time point, 50 µL of serum-free media and 50 µL of MTT solution (5 mg/mL in phosphate buffered saline) was added into each well followed by 4 hours incubation at 37°C. Addition of 150 µL of solvent (4 mM HCl, 0.1% NP40 in isopropanol) was followed by a 15 min incubation carried out in darkness with agitation at 4°C. Optical density was measured at 590 nm using a spectrophotometer.

### Quantitative RT-PCR

Total RNA was extracted from C28I2 cells using Trizol® (Invitrogen) following the manufacturer’s instructions. cDNA was prepared using 1 μg RNA with the High-Capacity cDNA Reverse Transcription Kit (Applied Biosystems). SYBR green based qPCR was performed in triplicate using human primers to Aggrecan (ACAN), Collagen 2 (Col2), NOS2, MMP13, ADAMTS5, and GAPDH (Applied Biosystems Real-time PCR system). mRNA levels were normalized to GAPDH and reported as relative mRNA fold change. Human primer sequences were as follows: ACAN: Forward: 5’ – AGGCAGCGTGATCCTTACC - 3’, Reverse: 5’ – GGCCTCTCCAGTCTCATTCTC - 3’; Col2: Forward: 5’- CCTGGCAAAGATGGTGAGACAG – 3’, Reverse: 5’ - GGCCTGGATAACCTCTGTGA – 3’; MMP13: Forward: 5’ – TGACCCTTGGTTATCCCTTG– 3’, Reverse: 5’ –ATACGGTTGGGAAGTTCTGG– 3’; NOS2: Forward: 5’ – ACCTCAGCAAGCAGCAGAAT – 3’, Reverse: 5’ – TGGCCTTATGGTGAAGTGTG – 3’; ADAMTS5: Forward: 5’ – CTTCACTGTGGCTCACGAAA – 3’, Reverse: 5’ – TTTGGACCAGGGCTTAGATG – 3’; GAPDH: Forward: 5’ - CGCTCTCTGCTCCTCCTGTT -3’, Reverse: 5’ - CCATGGTGTCTGAGCGATGT - 3’. Results were analyzed using the 2−ΔΔCq method.

### In vivo luminescent imaging

Atsttrin and E_5_C were labeled with Vivo-Tag 680-S in accordance with the manufacturer’s protocol excepting that conjugation was carried out overnight at 4°C. Free dye was removed using a 5 min centrifugation at 4000 rpm in a 3kDa MWCO Amicon filter. Adult DBA1/J mice, approximately 4 months of age, were anesthetized and hind limbs were shaved. 10 μL of phosphate buffered saline, Vivo-Tag 680-S labeled-E_5_C hydrogel, Vivo-Tag 680-S-labeled Atsttrin in E_5_C hydrogel, or Vivo-Tag 680-S-labeled Atsttrin in phosphate buffered saline were intra-articularly injected. Two mice were implemented in each group and injections were performed bilaterally to generate 4 regions of interest. Mice were imaged using PerkinElmer IVIS Spectrum imaging system (*PerkinElmer, MA, USA*) at days 0, 28 and 32 days (excitation 675 nm/ emission 725 nm). The fluorescence signal (p/s/cm2/sr) was normalized to background and measured using Aura imaging software (Spectral Instruments Imaging).

### Establishment of the Rabbit ACLT Model

All animal work was performed in accordance with protocols approved by the New York University IACUC. Animals were housed in a barrier facility maintained by New York University with *ad libitum* access to food and water. Skeletally mature male New Zealand rabbits (Charles River Laboratories) were subjected to either bilateral anterior cruciate ligament transection (ACLT, n=36) or a sham operation (n=6) under sterile conditions using a medial parapatellar approach whereby the patella was laterally dislocated and flexed maximally to reveal and identify the anterior cruciate ligament, which was transected using a sterile No. 11 blade [40, 53]. The knee joint was washed with sterile saline and closed using 4-0 nylon sutures and the skin incision was closed using 3-0 absorbable sutures. Appropriate transection of the ACL was confirmed via laxity of the cruciate observed via the anterior drawer and Lachman tests. Sham operated animals underwent the same procedure excluding transection of the ligament. For assessing the preventative potential of our biologics, intra-articular injection of 0.5 mL E_5_C alone (n=6), Atsttrin alone (0.5 mg) (n=6), or E_5_C loaded with Atsttrin (0.5mg) (n=6), or vehicle (sterile phosphate buffered saline; n=6) was performed intraoperatively. Animals were sacrificed 8 weeks after model induction. For animals in therapeutic studies, bilateral ACLT was performed followed by intra-articular injection of 0.5 mL E_5_C alone (n=3), E_5_C loaded with Atsttrin (0.5 mg; n=3), Atsttrin alone (0.5mg; n=3), or vehicle (n=3) after development of an osteoarthritic phenotype at 8 weeks post-ACLT [2]. E_5_C protein concentration was maintained at 1 mM in each group.

### Magnetic Resonance Imaging

Left limbs were harvested at sacrifice and immediately stored at 4°C prior to acquisition of magnetic resonance (MR) images within 36 hours of sacrifice. Scans were performed on a high field animal 7-T Bruker Biospec 70/30 available at the NYU Langone Preclinical Imaging Laboratory. The image acquisition protocol was optimized for the rabbit ACLT model. Imaging parameters were as follows: (1) 3-D spoiled gradient-recalled echo (SPGR) with fat suppression, echo time 5 ms, repetition time 20 ms, and flip angle 15°. In-plane matrix 320 x 320 with field of view (FOV) at 40x 40 giving 0.125 × 0.125 in-plane resolution, using 5 signal average amounting to 10 min 40 s total acquisition time. (2) For multi-slice, multi-echo (MSME)-T2 maps, with fat suppression: echo time, 8 ms; repetition time, 300 ms; and 25 echoes; slice thickness, 25 mm; section spacing, 0 mm; MRI field of view (FOV), 40 x 40 mm; matrix, 128 × 128 resolution, 0.31x 0.31 amounting to 10 min 14 s total acquisition time. All MR images were acquired sequentially in a coronal orientation. DICOM MR images were visualized in series using Analyze 14.0 (AnalyzeDirect). The patella, posterior cruciate ligament, medial collateral ligament, fibular collateral ligament attachment and meniscal horns were used as landmarks for anterior-posterior positioning and laterality. T2 weighted image stacks, with echo times between 10 and 80 ms, were imported into ImageJ and scaled for 100 um/pixel resolution using bicubic interpolation. T2 maps were generated via implementation of the Levenber-Marquardt fit algorithm (ImageJ MRI Processor plugin). Regions of interest highlighting weight bearing cartilage were drawn on the contrast enhanced T2 weighted image stack and imported to the T2 map for measurement in ImageJ.

### Macroscopic Evaluation of Articular Cartilage

Immediately following MRI acquisition, left hind limbs were disjointed. Fixed tissues were stained with India Ink for macroscopic evaluation of cartilage in accordance with a four-point scoring system wherein a score of 1 reflected an intact cartilage, 2 indicated minimal fibrillation, 3 signaled overt fibrillation and a score of 4 reflected full thickness erosion [42].

### Micro-CT

Micro-CT scanning of right fixed femurs was completed on a Skyscan 1172 (Bruker Corporation, USA) with parameters as follows: 8 µm isotropic voxel size, 80 kV, 124 mA, 975 ms exposure, 0.600° rotation step, frame averaging value 10, 0.5 mm Aluminum filter. Reconstructed axial datasets were implemented for assessment of subchondral bone quality using analysis toolkits in CT-Analyzer (Bruker). Volumes of interest were selected distal to the appearance of growth plate bone bridging and extended approximately 6 mm to exclude the subchondral bone plate. After global thresholding for selected volumes, three circular (2 mm diameter) regions of interest were placed within the anterior, central, and posterior aspects of the weight-bearing region of the medial and lateral femoral condyles to generate 6 regions of interest for each animal (n=3/group) [45]. 3D analysis was performed to assess bone volume fraction (BV/T, %), trabecular thickness (Tb.Th; mm), trabecular number (Tb.N; mm^−1^), and trabecular spacing (Tb.S; mm) in each region. Bone mineral density (BMD; g.cm^−3^) phantoms were implemented for calibration of BMD calculations. Osteophyte volume was assessed by manual segmentation of osteophytic tissue based on deviation from the appearance of the underlying normal bone contour; manual segmentation was performed every 5 slices with adaptive interpolation over the entirety of the image sequence (approximately 10 mm). The total 3D tissue volume of this region of interest was used to quantify osteophyte volume. All analyses were performed following application of a Gaussian filter, despeckle function, and global thresholding.

### Histopathological examination of joints

Following routine fixation, decalcification, and paraffin embedding, 6 µm serial sections were prepared and stained with 0.1% Safranin O/Fast Green and Movat Pentachrome. Safranin O-stained sections were used to grade the extent of cartilage destruction in accordance with OARSI scoring guidance for rabbit ACLT. Briefly, tissues were scored from 0 to 6 across Safranin O–fast green staining with a score of 0 reflecting uniform staining and a score of 6 indicating loss of staining in all the hyaline cartilage ≥50% the length of the condyle or plateau, cartilage structure was scored on an 11 point scale with 0 reflecting intact surface and 11 indicating full depth erosion hyaline and calcified cartilage to the subchondral bone ≥50% surface, chondrocyte density was assessed on a 4 point scale with 0 indicating no reduction in cell density and 4 indicating global reduction in cell density, and cluster formation was scored on a 3 point scale with 0 assigned for the absence of cluster formation and a score of 3 indicating the presence of ≥8 clusters. Score was assessed in four compartments: medial femur, lateral femur, medial tibia and lateral tibia. Minimal and maximal possible scores for each compartment were 24. Total joint score for each limb was taken by summation of scores for the four compartments.

### Immunohistochemistry

For immunohistochemistry of indicated biomarkers, deparaffinized and rehydrated sections were first subjected to sequential enzymatic digestion with 250 U/mL hydraulinase, 100 mU/mL chondroitinase ABC, and 0.1% trypsin for 30 min at 37℃. Sections were blocked with 5% normal goat serum for 60 min at room temperature. Tissues were incubated with anti-Aggrecan Neoeptiope (NITGE) (1:50, MBS442004, Mybiosource) or anti-MMP13 (1:25, MA5-14238, ThermoFisher) primary antibodies. Sections were then washed thrice in phosphate buffered saline prior to 30 min incubation with biotin-conjugated Goat Anti-Mouse IgG (H+L) (1:500, BP-9200, Vector Laboratories). Endogenous peroxidase was blocked and detection was performed using VECTASTAIN® ABC-HRP Kit with signal development carried out using 3,3-diaminobenzidine. Slides were counterstained with 0.5% Methyl green.

### Statistical analyses

All data are presented as means ± SEM. The numbers of animals used per group are indicated in figure legends. ANOVA with multiple comparisons and post hoc Tukey test was used when comparing multiple groups. A value of P < 0.05 was considered statistically significant.

## Acknowledgment

This work is supported partly by NIH research grants R01AR062207, R01AR061484, R01AR076900, R01NS103931, and a DOD research grant W81XWH-16-1-0482. The NYU Microscopy Core is partially supported by NYU Cancer Center Support Grant NIH/NCI P30CA016087. SEM is supported by the MRI program of the National Science Foundation under Award DMR-0923251.The authors would like to acknowledge all lab members for insightful discussions. We also thank NYU Dentistry microCT core for technical support.

## Author contributions

P.K., A.H., R.R., J.K.M. and C-J.L. designed the experimental plan and wrote the manuscript. P.K. and A.H. planned and executed the experiments. M.M. and B.L. assisted in production and characterization of E_5_C construct. C.C, M.C. and G.S performed the surgical OA model. S.H. performed and analyzed rtPCR experiments. R.M. performed MR imaging. R.R. and R.M. provided consultation on MR data analysis.

