## Supplemental figures for "Injectable Progranulin-derivative Atsttrin loaded Protein Engineered Gels for Post Traumatic Osteoarthritis"

**Supplementary Information**

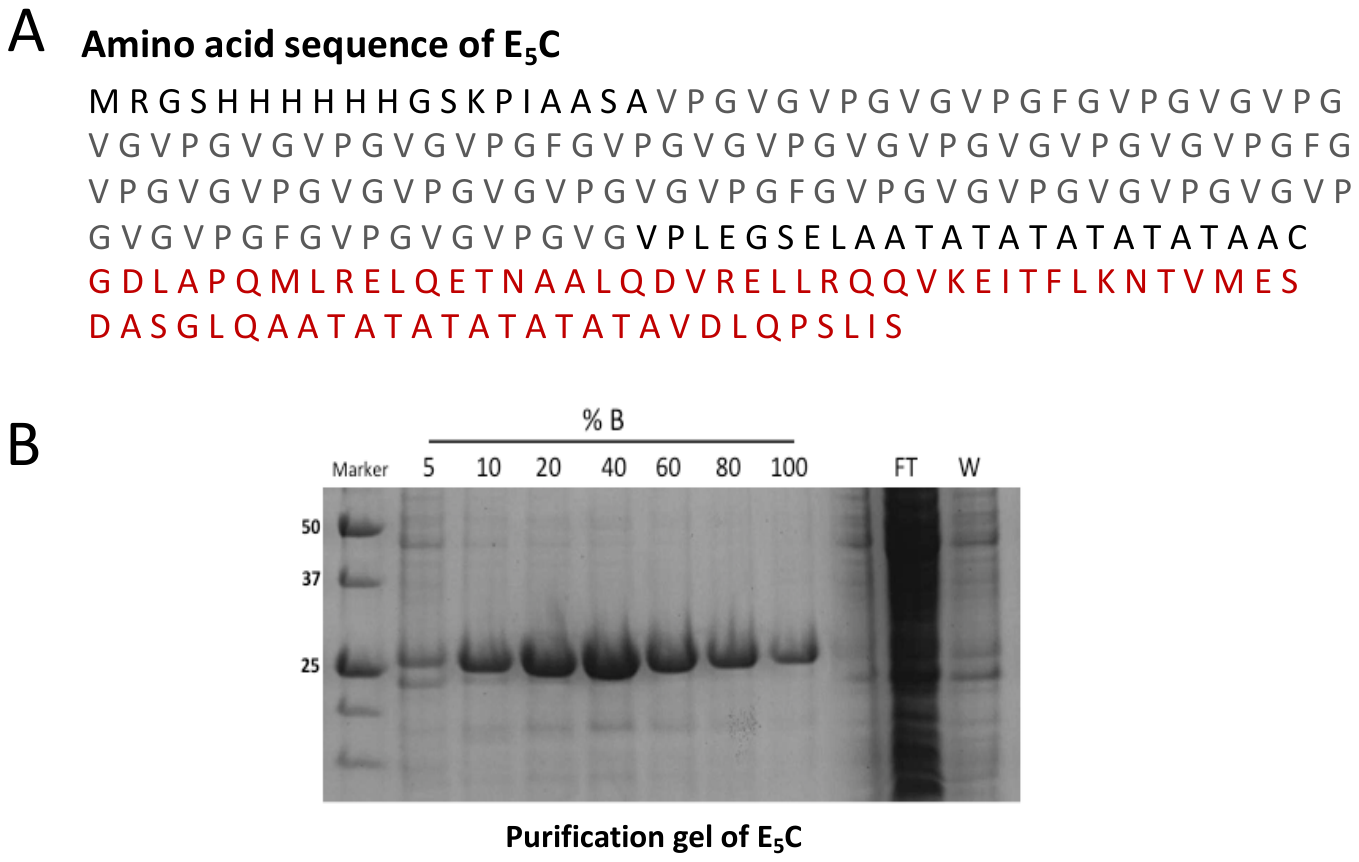

**Fig. S1.** (A) Amino acid sequence of E_5_C. Five repeats of E are indicated in gray while C domain is indicated in red.(B) SDS-PAGE analysis of different fractions eluted using a concentration gradient of imidazole ranging from 5-100%. *FT: Flow-through, W=Wash.*

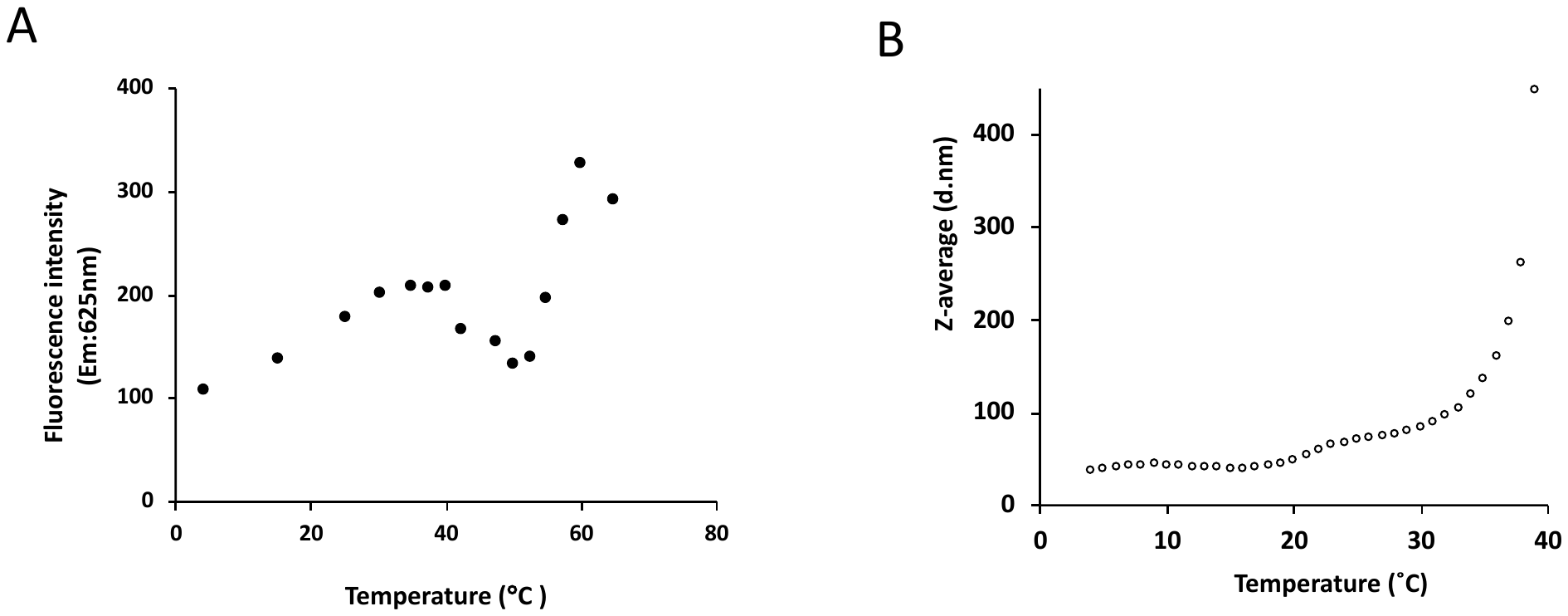

**Fig. S2.** Critical Micelle Temperature studies using Nile Red probe fluorescence. (B) Z-average diameter of E_5_C indicating a gradual increase in particle size with a drastic increase in size beyond physiological temperature.

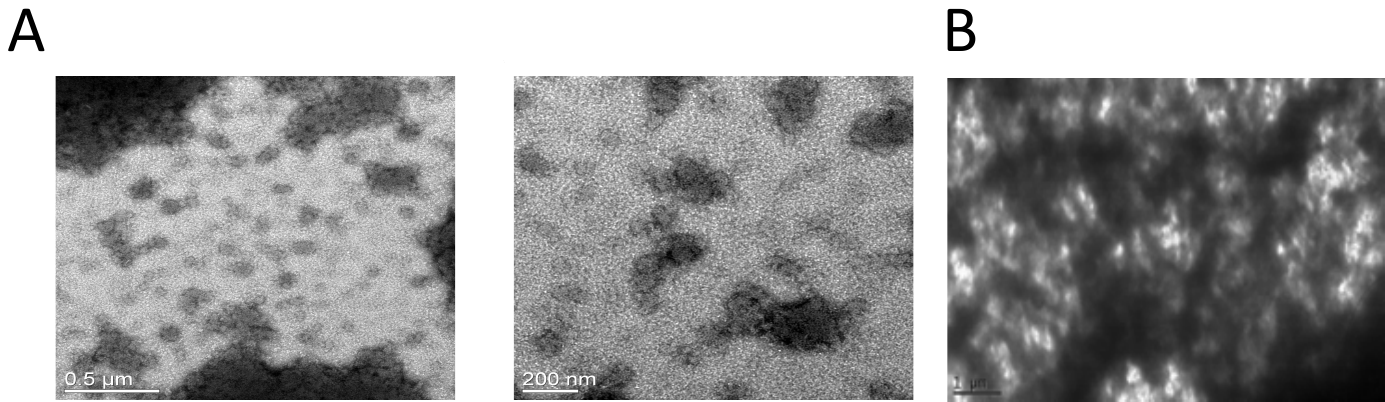

**Figure S3**. Transmission electron Micrographs showing (A) micelles formation and (B) micelle packing that lead to network formation of E_5_C.
